# Tagsteady: a metabarcoding library preparation protocol to avoid false assignment of sequences to samples

**DOI:** 10.1101/2020.01.22.915009

**Authors:** Christian Carøe, Kristine Bohmann

## Abstract

Metabarcoding of environmental DNA (eDNA) and DNA extracted from bulk specimen samples is a powerful tool in studies of biodiversity, diet and ecological interactions as its inherent labelling of amplicons allows sequencing of taxonomically informative genetic markers from many samples in parallel. However, the occurrence of so-called ‘tag-jumps’ can cause incorrect assignment of sequences to samples and artificially inflate diversity. Two steps during library preparation of pools of 5’ nucleotide-tagged amplicons have been suggested to cause tag-jumps; i) T4 DNA polymerase blunt-ending in the end-repair step and ii) post-ligation PCR amplification of amplicon libraries. The discovery of tag-jumps has led to recommendations to only carry out metabarcoding PCR amplifications with primers carrying twin-tags to ensure that tag-jumps cannot result in false assignments of sequences to samples. As this increases both cost and workload, a metabarcoding library preparation protocol which circumvents the two steps that causes tag-jumps is needed. Here, we demonstrate Tagsteady, a metabarcoding Illumina library preparation protocol for pools of nucleotide-tagged amplicons that enables efficient and cost-effective generation of metabarcoding data with virtually no tag-jumps. We use pools of twin-tagged amplicons to investigate the effect of T4 DNA polymerase blunt-ending and post-ligation PCR on the occurrence of tag-jumps. We demonstrate that both blunt-ending and post-ligation PCR, alone or together, can result in detrimental amounts of tag-jumps (here, up to ca. 49% of total sequences), while leaving both steps out (the Tagsteady protocol) results in amounts of sequences carrying new combinations of used tags (tag-jumps) comparable to background contamination.

## Introduction

Metabarcoding of DNA extracted from environmental and community samples has become a valuable tool in studies on biodiversity, diet and ecological interactions (e.g. Taberlet et al. 2012; Bohmann et al. 2014; Alberdi et al. 2018). Metabarcoding relies on PCR amplification with metabarcoding primers targeting a taxonomically informative marker within a selected taxonomic group (Taberlet et al. 2012). The basis of metabarcoding is the addition of sample-specific nucleotide identifiers to amplicons and the use of these to assign metabarcoding sequences back to the samples from which they originated. An often used approach to achieve this is a tagged PCR approach in which sample DNA template is amplified with metabarcoding primers carrying 5’ nucleotide tags of typically 6-10 nucleotides in length (first demonstrated by Binladen et al. 2007). Subsequently, differently tagged amplicons are pooled and built into sequencing libraries for the chosen sequencing platform. This allows for the taxonomic content of many samples to be sequenced in parallel on high-throughput sequencing platforms (e.g. Valentini et al. 2009; Bohmann et al. 2011; Shehzad et al. 2012).

The effectiveness of metabarcoding relies on the ability to correctly track tagged amplicons back to the samples from which they originated. However, amplicon sequences carrying false combinations of used nucleotide tags, so-called tag-jumps, have been reported both on the Roche/454 sequencing platform (Rakel Blaalid et al. 2013; Carew et al. 2013; Lindner et al. 2013; Davey et al. 2013; Davey, Kauserud, and Ohlson 2014; Botnen et al. 2014; R. Blaalid et al. 2014) and on the Illumina sequencing platform (Esling, Lejzerowicz, and Pawlowski 2015; Schnell, Bohmann, and Gilbert 2015) where up to 28.2% misassigned unique sequences have been identified (Esling, Lejzerowicz, and Pawlowski 2015). Such tag-jumps may introduce significant levels of incorrect assignments of sequences to samples, leading to false positives and artificial inflation of diversity in the samples, much to the detriment of metabarcoding studies (Schnell, Bohmann, and Gilbert 2015; Esling, Lejzerowicz, and Pawlowski 2015). Tag-jumps can be accounted for in the experimental setup through twin-tagging of amplicons, i.e. using forward and reverse primers carrying matching tags, F1-R1, F2-R2, etc. (Schnell, Bohmann, and Gilbert 2015). However, this greatly increases both cost and workload of metabarcoding studies as a higher number of differently tagged primers and library preparations are needed, which ultimately restricts sample throughput (Schnell, Bohmann, and Gilbert 2015). Further, although tag-jumps can be identified through such a setup, a proportion of the sequencing output will be wasted on the reads carrying tag jumps. Thereby, the tag-jumps ultimately adds to the sequencing cost of correctly assigned reads.

Tag-jumps have been suggested to occur in two steps during library preparation of pools of tagged amplicons: i) when using T4 DNA Polymerase for blunt-ending prior to ligation of sequencing adapters and ii) as a consequence of chimera formation during post-ligation PCR (van Orsouw et al., 2007; Esling et al., 2015; Schnell et al., 2015; Palkopoulou et al., 2016). The T4 DNA Polymerase is a 3’→5’ exonuclease possessive enzyme (Rittié and Perbal 2008). When tagged amplicons are pooled prior to library preparation, semi-complementary single strands of amplicons carrying different tags can hybridize along the complementary parts (the metabarcoding marker and primer regions), keeping the terminal nucleotide tags as single-stranded overhangs (Figure 1). Of these, the 3’ overhangs become substrate for the 3’→5’ exonuclease activity of T4 DNA Polymerase, and the enzyme will degrade the un-matching nucleotide tag at the 3’ end. The opposite strand, the 5’ overhangs (i.e. the inherent tag), will then act as a template for extension, causing the tag to ‘jump’ from one strand to the other (van Orsouw et al. 2007; Schnell, Bohmann, and Gilbert 2015). Van Orsouw et al. (2007) reduced the occurrence of tag-jumps by using tagged primers with a 5’-phosphate and omitting the blunt-ending step during library preparation for the 454 FLX platform. The effect of completely excluding post-ligation PCR amplification of pools of tagged amplicons has to our knowledge not been investigated.

**Figure 1.**
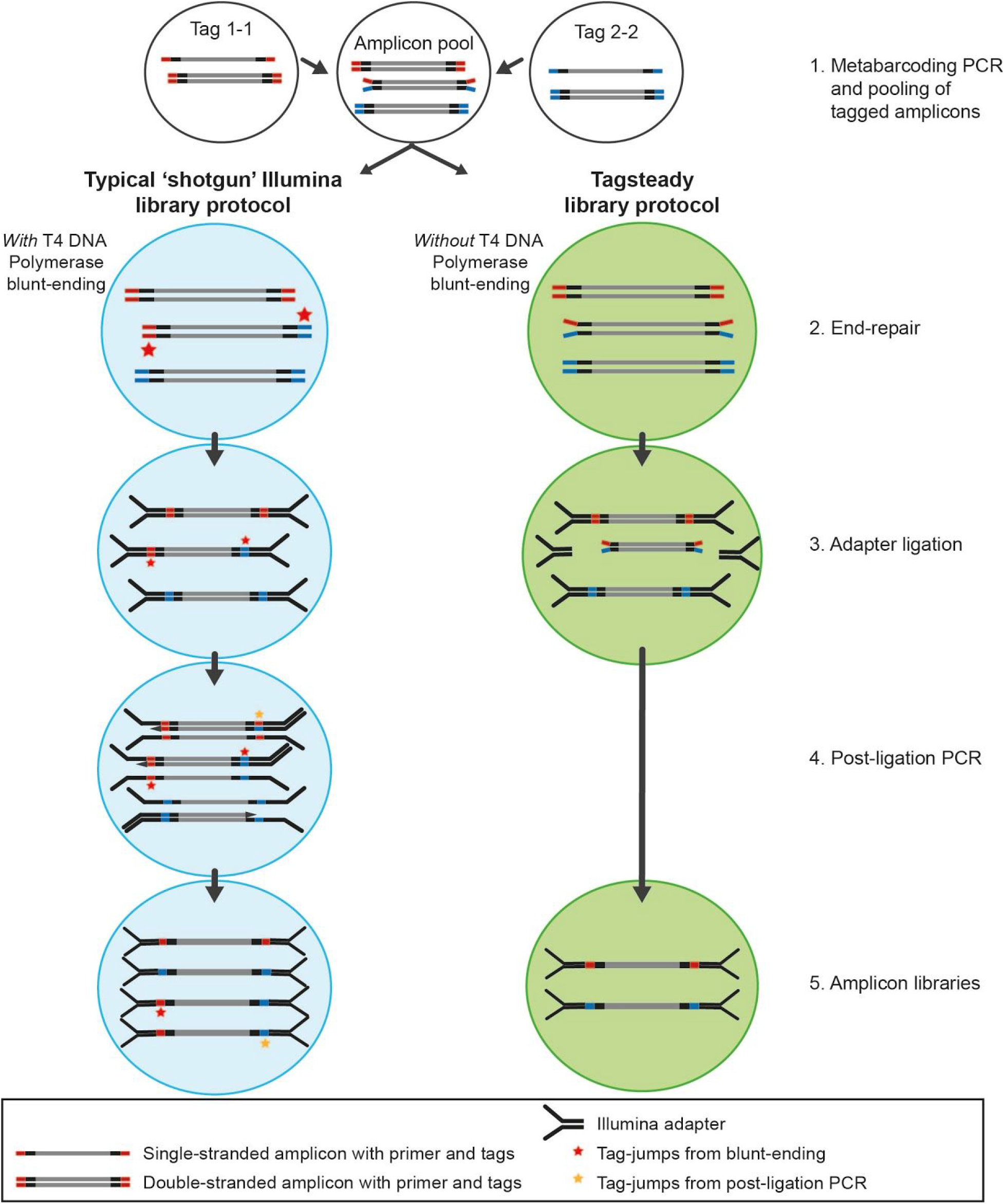
Overview of metabarcoding and library preparation steps and formation of tag-jumps in a typical ‘shotgun’ Illumina library protocol and our presented Tagsteady library protocol. 1) Metabarcoding PCR with 5’ nucleotide tagged primers. To allow detection of tag-jumps, only unique twin-tag combinations is visualised. Following pooling of PCR reactions, differently tagged single-stranded amplicons can form heteroduplexes with non-complementary tag overhangs. 2) In a typical ‘shotgun’ Illumina library protocol (left), T4 DNA polymerase is used for blunt-ending, T4 PNK for 5’ phosphorylation and Taq polymerase for 3’ adenylation. In this type of end-repair, 3’ overhangs (in heteroduplexes) will become substrate for the 3’→5’ exonuclease activity of T4 DNA Polymerase. The opposite strand, the 5’ overhangs (i.e. the inherent tag), will then act as a template for extension, causing the tag to ‘jump’ from one strand to the other (asterisk) (see van Orsouw et al. 2007; Schnell, Bohmann, and Gilbert 2015). The Tagsteady end-repair (right) only contains T4 PNK and Klenow exo- (thus no exonuclease activity) and therefore tag-jumps cannot arise. 3) After end repair, T4 DNA Ligase is used to ligate Illumina sequencing adapters (here depicted as Illumina Y-shaped adapters). 4) Often post-ligation PCR is carried out, causing further tag-jumps as a result of incomplete primer extension. Post-ligation PCR is not necessary with the Tagsteady protocol as it uses PCR-free full length adapters. 5) Sequencing of libraries on an Illumina sequencing platform. 6) Following initial sequence read processing, sequences within each amplicon library are sorted according to primer and tag sequences to assess levels of sequences carrying new combinations of used tags (tag-jumps).

Today, the most popular high-throughput sequencing platform for metabarcoding is the Illumina series due to its wide availability, high-throughput, and relatively low error rates and long paired-end reads (www.illumina.com, applied in e.g. Shehzad et al. 2012; Quéméré et al. 2013; Hope et al. 2014; Elbrecht et al. 2017; Stoeck et al. 2018; Singer et al. 2019). PCR-free library preparation protocols that circumvent the post-ligation PCR enrichment step, and thereby between-sample chimeras, have been developed for Illumina sequencing (e.g. Kozarewa et al. 2009) and embodied in commercial kits, such as Illumina TruSeq (applied in e.g. Apothéloz-Perret-Gentil et al. 2017). However, these protocols or commercial kits were originally designed for library preparation for shotgun sequencing and therefore include a blunt-ending step to repair fragmented DNA with overhangs in preparation for adapter ligation (e.g. Kozarewa et al. 2009; Neiman et al. 2012; Carøe et al. 2018; Zheng et al. 2010; Bentley et al. 2008; Margulies et al. 2005). Therefore, while these protocols will eliminate tag-jumps caused by chimera formation between tagged amplicons during library-PCR amplification, they will not eliminate those caused by T4 DNA Polymerase or similar activity (van Orsouw et al. 2007; Palkopoulou et al. 2016). The current recommendation for metabarcoding with nucleotide-tagged primers is to account for tag-jumps by only using primers carrying twin-tags to ensure that tag-jumps cannot result in false assignments of sequences to samples. As mentioned above, this greatly increases both cost and workload of metabarcoding studies, which ultimately restricts sample throughput (Schnell, Bohmann, and Gilbert 2015). There is therefore a great need for a metabarcoding library preparation protocol, which circumvents the two steps that cause tag-jumps.

In this study, we present and validate ‘Tagsteady’, a virtually tag-jump free library preparation protocol for Illumina sequencing of pools of tagged amplicons. The Tagsteady protocol is developed as a single-tube library preparation protocol, circumventing both the use of T4 DNA Polymerase in the end-repair step and the post-ligation PCR amplification step. To document the functioning of the Tagsteady protocol and assess how T4 DNA Polymerase blunt-ending and post-ligation PCR affect the prevalence of tag-jumps, we carried out four library preparation protocol treatments on six pools of twin-tagged amplicons. The four library protocol treatments represent combinations with and without T4 DNA Polymerase blunt-ending and with and without post-ligation PCR (Figure 2). Moreover, to assess whether the Tagsteady protocol can withstand high proportions of single-stranded amplicons, we applied it to a set of denatured and re-hybridised amplicon pools. Finally, to validate the robustness and stability of the Tagsteady protocol, we use it to build libraries of a further 15 pools of twin-tagged amplicons (Figure 2).

**Figure 2.**
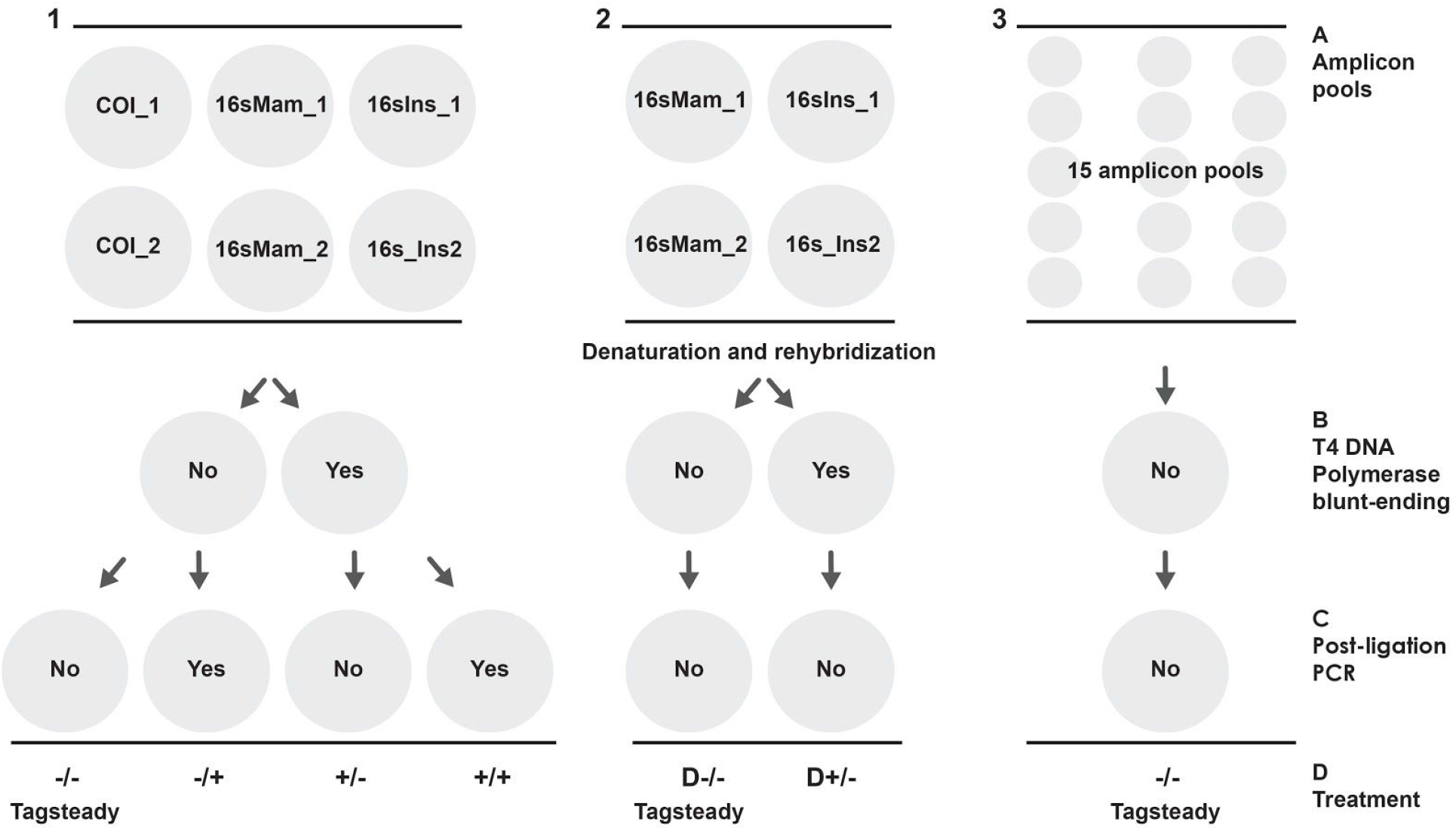
Experimental overview. 1A) Six pools of 5’ twin-tagged amplicons generated with three metabarcoding primer sets were used to assess the effect of blunt-ending and post-ligation PCR on tag-jumps. Each of the six amplicon pools were subjected to four different treatments. 1B) The four treatments represent combinations with and without T4 DNA Polymerase blunt-ending in the end-repair step and 1C) with and without post-ligation PCR. 1D) This resulted in 24 libraries for the 6 amplicon pools, representing four library preparation treatments for each amplicon pool. 2A) To further assess the effect of T4 DNA Polymerase blunt-ending on the prevalence of tag-jumps, we denatured and re-hybridised four amplicon pools (16sMam1/2 and 16sIns1/2). 2B) End-repair was carried out with and without T4 DNA Polymerase blunt-ending and with no post-ligation PCR (2C). 3) Finally, to validate the robustness and stability of the Tagsteady protocol, we applied it to 15 pools of twin-tagged amplicons.

## Materials and methods

### Design of end-repair and adapter ligation steps

To assess the effect of blunt-ending, we formulated two PCR-free library protocols, mainly differing in the addition of T4 DNA Polymerase and its ability to digest 3’ overhangs and perform blunt-ending. The two protocols only differed in the reagents and enzymes found in the end-repair reaction, i.e. they had identical ligation steps (Figure 1).

The first protocol (hereafter referred to as ‘+’) included a blunt-ending step in the end-repair (Figure 1). Normally, end-repair for fragmented DNA for PCR-free Illumina library building is dependent on three enzymatic activities, namely 5’ phosphorylation, blunt-ending (by 3’ overhang digestion and 5’ overhang fill-in) and 3’ adenylation (Kozarewa et al. 2009). Similar to recent protocols, we formulated this as a so-called ‘single-tube’ protocol, in which T4 Polynucleotide Kinase (T4 PNK) is used for 5’ phosphorylation, T4 DNA Polymerase for blunt-ending (filling in 3’ recessed ends and degrading 3’ overhangs) and finally Taq DNA Polymerase for 3’ adenylation through its deoxynucleotidyl transferase activity. The single-tube design relies on using heat-inactivation of the enzymes rather than purification prior to the ligation of sequencing adapters (Rittié and Perbal 2008; Carøe et al. 2018; Neiman et al. 2012).

We designed the second protocol (hereafter referred to as ‘−’) explicitly for library preparation of 5’ tagged amplicons and to test the effect of omission of T4 DNA Polymerase blunt-ending on the prevalence of tag-jumps (Figure 1). In the development of the end-repair step in the second protocol, we took advantage of the fact that amplicons are double-stranded DNA molecules that either have blunt ends or when using a Taq polymerase for the PCR, possess a 3’ adenine overhang (Rittié and Perbal 2008). As we, and many others, use Taq DNA polymerase in our metabarcoding PCR amplifications, we mainly based the Tagsteady end-repair on the 5’ phosphorylation activity of T4 PNK. However, to further enable 3’ adenylation of partially adenylated amplicons or blunt-end amplicons (depending on the type of polymerase used in the metabarcoding PCR), we also included Klenow Fragment 3’→5’ exo- and dATP in the Tagsteady end-repair step. We designed the Tagsteady protocol as a single-tube protocol by replacing purification with heat-inactivation prior to adapter ligation.

To circumvent the post-ligation PCR (often needed for enrichment and to index the libraries with indexed primers, see e.g. Meyer & Kircher (2010), we designed Illumina full-length Y-adapters (Kozarewa et al. 2009) with 3’ dT overhangs (facing insert) and dual matching indices, i.e. p5-p7: 1-1, 2-2, etc. to ensure that library bleeding on the flow-cell did not cause false assignment of sequences to libraries (Sinha et al. 2017; Kircher, Sawyer, and Meyer 2012). The indices were 7 nucleotides in length with a minimum of three mismatches between them (Supplementary information S1).

### Experimental design

To validate the Tagsteady protocol and assess the effect of T4 DNA Polymerase blunt-ending and post-ligation PCR on the prevalence of tag-jumps, we performed library preparation on six pools of twin-tagged amplicons with four different treatments representing combinations of the two PCR-free library preparation protocols mentioned above and with and without post-ligation PCR; i) a library protocol with blunt-ending and *with* a post-ligation PCR (referred to as ‘+/+’), ii) no blunt-ending (−) and *with* post-ligation PCR (referred to as ‘−/+’), iii) with blunt-ending and *no* post-ligation PCR (referred to as ‘+/−’), and iv) the Tagsteady protocol, i.e. no blunt-ending (−) and no post-ligation PCR (referred to as −/−) (Fig. 2; Supplementary Information Table S2). Further, to assess whether the Tagsteady protocol can withstand high levels of single-stranded amplicons generated in the metabarcoding PCR, we denatured and re-hybridized aliquots of four of the amplicon pools and subjecting them to the Tagsteady protocol (−/−) and the ‘+/−’ library protocol with blunt-ending and *no* post-ligation PCR. These are denoted D−/− and D+/−, respectively, with the D indicating the denaturing treatment. Finally, we validated the Tagsteady protocol (−/−) on 15 pools of twin-tagged amplicons.

### Pools of tagged amplicons

We used six pools of 5’ nucleotide-tagged amplicons from two metabarcoding studies: metabarcoding of metazoan and vertebrate DNA in vampire bat gut contents (Bohmann et al. 2018) and of insect DNA in bat faecal extracts (unpublished). Importantly, each PCR amplification was carried out with twin-tags (F1-R1, F2-R2, etc.) to ensure tag-jump events could be identified. The six pools of tagged amplicons used in this study included: i) two pools of tagged amplicons from vampire bat gut contents, consisting of amplicons generated through PCR amplification with mammal mitochondrial 16s rRNA primers (16smam1/16smam2, from here on referred to as ‘16sMam’), amplifying a ca. 95 bp fragment excluding primers (Taylor 1996), ii) two pools of tagged amplicons from vampire bat gut contents, PCR amplified with metazoan COI primers (mlCOIintF/jgHCO2198, from here on referred to as ‘COI’), amplifying 313 bp fragment of the COI barcode region excluding primers (Geller et al. 2013; Leray et al. 2013; Bohmann et al. 2018), and iii) two pools of tagged amplicons from insect-eating bat faecal droppings, PCR amplified with insect mitochondrial 16sRNA primers (Ins16s_1shortF /Ins16s_1shortR, from here on referred to as ‘16sIns’), amplifying a ca. 190 bp fragment (excluding primers) (Clarke et al. 2014; Elbrecht et al. 2016).

Metabarcoding primers were tagged by adding nucleotides at the 5’ ends (Binladen et al. 2007). The 16sMam primers carried 5’ nucleotide tags that were 7-8 bp long (see Bohmann et al. 2018). The COI and 16sIns primers carried 5’ nucleotide tags of 7-8 nucleotides in length of which 6 nucleotides were the actual tag and 1-2 nucleotides were added to increase complexity on the flow-cell (De Barba et al. 2014). For the 16sIns, these additional nucleotides were specified, while for the COI they were ordered as random nucleotides. Tags were designed to have a minimum of 3 nucleotide mismatches between any pair of tags. Each of the three primer sets consisted of 60 tagged forward and 60 tagged reverse primers.

All PCRs were set up in a dedicated pre-PCR laboratory with positive air pressure to minimize risk of contamination. PCR amplifications and post-PCR laboratory steps, including library preparations, were carried out in a dedicated post-PCR laboratory. In all laboratories, handling of reagents, samples, DNA extracts, PCR products and sequence libraries was performed in laminar flow hoods.

Prior to metabarcoding PCRs, for all three primer sets quantitative PCR was carried out to optimise metabarcoding PCR conditions and assess negative extraction controls as described in Bohmann et al. (2018). The 16sIns PCRs were carried out in 25 μl reactions containing 1 μl template DNA, 1 U AmpliTaq Gold, 1x Gold PCR Buffer and 2.5 mM MgCl_2_ (all from Applied Biosystems), 0.2 mM dNTP mix (Invitrogen), 0.5 mg/ml BSA, 0.6 μM of each 5’ nucleotide-tagged forward and reverse primer. PCR conditions for 16sIns were as follows: 95°C for 10 minutes, 35 cycles of 95°C for 15 seconds, 54°C for 30 seconds and 72°C for 30 seconds and a seven minutes final extension at 72°C. PCR setup and conditions for the 16sMam and COI primer sets can be found in Bohmann et al. (2018). For all three primer sets, a negative PCR control was included for every seven samples. PCR products were visualised on 2% agarose gels. Negative controls did not show identifiable amplification. PCR products were pooled at approximately equimolar ratios determined by gel band strength. Negative controls were added in the same volume as sample extracts showing the weakest bands. The six amplicon pools each contained 45-49 twin-tagged PCR products of which 38-44 were from samples, one from positive controls and 3-7 from negative controls. Following pooling, the amplicon pools were purified using 1.8x SPRI bead solution prepared following Rohland & Reich (2012), see Faircloth and Glenn (2014). Briefly, 1.8x SPRI bead solution was added to each amplicon pool, mixed by vortexing and incubated at room temperature for ten minutes. Subsequently, the beads were pelleted with a magnet and washed twice with 80% ethanol and air dried for five minutes before DNA was eluted in EB buffer (Qiagen). Purified amplicon pools were quantified using a Qubit Fluorometer (Invitrogen) using HS reagents. The amount of moles within each amplicon library was calculated based on Qubit measurements and amplicon length (incl. primers and nucleotide tags).

To mimic the effect of large amounts of single-stranded DNA generated in the metabarcoding PCR, we denatured aliquots of four of the aforementioned amplicon pools (two 16sMam and two 16sIns amplicon pools) and subsequently re-hybridized these to form double-stranded DNA. This was done in 50 μl reactions in 0.2 ml PCR tubes in a thermocycler in which we denatured for two minutes at 95°C followed by cooling down to room temperature over 20 minutes. Each of the four denatured pools were then subjected to the two library protocol treatments without post-ligation PCR; D−/− and D+/−.

To validate the stability of the Tagsteady protocol (−/−), we further used it to build libraries on 15 pools of twin-tagged amplicons generated with the insect mitochondrial 16sRNA primers (16sIns) mentioned above. The 15 amplicon pools each consisted of 34-49 twin-tagged PCR products of which 29-44 were from samples, 1-2 from positive controls and 3-4 from negative controls.

### Library preparation

For each library reaction, 2 pmol purified amplicon pool was adjusted to 30 μl with molecular grade water just prior to library build. In addition, for each of the four library protocol treatments, two negative library controls were included in which 30 μl molecular grade water replaced the amplicon template. During each step, reagents were kept on ice and reactions were set up on ice blocks. All library incubations were performed in 0.5 ml Lobind Eppendorf tubes in an Applied Biosystems 2720 thermal cycler.

End-repair without T4 DNA Polymerase: for treatments −/+ and −/−, an end-repair mastermix was made by combining 4 μl T4 DNA ligase reaction buffer (New England Biolabs, NEB, Ipswich, Massachusetts, US), 0.5 μl dATP (10mM) (Thermo-Fisher), 2 μl reaction booster mix (consisting of 25 % PEG-4000 (Sigma Aldrich, 50%), 2 mg/ml BSA (Thermo-Fisher) and 400 mM NaCl) (Mak et al. 2017), 2 μl T4 PNK (NEB, cat#M0201S, 10 U/μl) and 1.5 μl Klenow Fragment (3’->5’ exo-) (NEB, cat#M0212S, 5 U/μl) per amplicon pool reaction. Ten μl of this mastermix was then added to each 30 μl amplicon pool, mixed well by pipetting and incubated for 30 minutes at 37°C followed by 30 minutes at 65°C and finally cooled to 4°C. No purification of the end-repair reactions was performed prior to ligation.

End-repair with T4 DNA Polymerase: for the treatments +/− and +/+, an end-repair mastermix was made by combining 4 μl T4 DNA Ligase Reaction Buffer (NEB), 0.5 μl dNTP (10mM) (Thermo-Fisher), 2 μl reaction booster mix (25% PEG-4000 (Sigma Aldrich, 50%), 2 mg/ml BSA (Thermo-Fisher) and 400 mM NaCl), 2 μl T4 PNK (NEB, cat#M0201S, 10 U/μl), 1.5 μl T4 DNA Polymerase (NEB, cat#M0203S, 3 U/μl) and 0.1 μl Taq DNA Polymerase (NEB, cat#M0273S, 5 U/μl). 10.1 μl of this mastermix was added to each of the 30 μl amplicon pools and incubated for 30 minutes at 20°C followed by 30 minutes at 65°C and finally cooled to 4°C.

Adapter ligation: All four treatments included the following adapter ligation protocol: 2 μl (20 μM) dual-index p5-p7 adapter mix (Supplementary Information S1) was added to each end-repaired amplicon pool, reaching a total volume of 42 μl for treatments −/+ and −/− and 42.1 μl for +/− and +/+. Each adapter mix contained unique p5/p7 matching indices to account for potential bleeding on the flow cell. Reactions were mixed well by pipetting the entire volume 10 times. A ligation mastermix was made consisting of 1 μl T4 DNA Ligase (NEB, cat#M0202S, 400 U/μl), 1 μl of T4 DNA ligase buffer (10x stock) and 6 μl of PEG-4000 (Sigma Aldrich, 50%) per amplicon pool reaction. Eight μl of this ligation mastermix was then added to each reaction for a final reaction volume of 50 μl and mixed well by pipetting. Ligation reactions were incubated for 20 minutes at 20°C followed by 10 minutes at 65°C to heat inactivate the enzyme, before cooling to 4°C. Libraries were purified with 1x volume of SPRIbeads as previously described and eluted in 30 μl EB buffer (Qiagen).

### Post-ligation PCR

To assess the effect of post-ligation PCR amplification on the amount of tag-jumps, two of the library protocol treatments included PCR amplification using IS7 and IS8 primers targeting the terminal 3’ end of the Illumina adapters (Meyer and Kircher 2010). Each PCR was carried out in 50 μl reactions with 2.5 μl of library template and final concentrations of 1x AmpliTaq Gold buffer II, 2.5 mM MgCl_2_, and 0.04 U/μl Amplitaq Gold polymerase (Thermo-Fisher, cat#N8080241), 0.25 mM dNTP (Thermo-Fisher), 0.8 mg/ml BSA (Thermo-Fisher) and 0.4 μM forward/reverse primer. PCR conditions were 95°C for 10 minutes, followed by 12 cycles of 95°C for 30 seconds, 60°C for 40 seconds and 72°C for 60 seconds followed by a final extension of 7 minutes at 72°C. PCR amplifications were performed on an Applied Biosystems 2720 thermal cycler. Amplified libraries were purified with 1x volume of SPRIbeads as described above and eluted in 30 μl EB buffer (Qiagen).

### Quantification of libraries and sequencing

Indexed libraries were quantified using quantitative PCR, qPCR, on an Agilent mx3005p instrument using New England Biolabs NEBNext^®^ Library Quant Kit for Illumina (NEB, cat#E7630S). For each amplicon library, quantifications were done on two 1:10,000 library dilution replicates, each consisting of 10 μl reactions with 2 μl template and otherwise run according to the manufacturer’s protocol. Indexed amplicon libraries originating from each of the three metabarcoding primer sets (16sMam, COI and Ins16s) were pooled together in equimolar ratio resulting in three sequencing pools. Further, the 15 16sIns amplicon pools built into libraries with the Tagsteady protocol were pooled together. The four sequencing pools were spiked with PhiX to increase complexity on the flow-cell and sequenced on an Illumina MiSeq instrument in paired-end mode running 150-250 cycles depending on the insert size of the sequence library. Sequencing was carried out by the Danish National High-Throughput Sequencing Center, University of Copenhagen, Denmark.

### Data analysis

Using AdapterRemoval version 2.2.0 (Schubert, Lindgreen, and Orlando 2016), sequence reads were trimmed for adapters, consecutive stretches of Ns and low-quality bases (minquality 28) and only sequences with a minimum length of 50 bp for the 16sMam and 16sIns amplicon libraries and 150 bp for the COI amplicon libraries were retained. Further, paired-end reads were merged with a minimum alignment length of 20 bp for the 16sMam and 16sIns amplicon libraries and 50 bp for the COI amplicon libraries. Using a modified version of DAMe (https://github.com/shyamsg/DAMe, Zepeda-Mendoza et al. 2016), sequences within each amplicon library were sorted according to primer and tags, keeping only sequences with perfect matches to forward and reverse primer sequences and tag sequences. Within each amplicon library, sequences were assigned to four categories based on the nucleotide tags they carried: i) tag combinations where the tag pair was used in the amplicon pool, ii) tag combinations where both tags were used in the amplicon pool, but not in this combination (tag-jumps), iii) tag combinations where only one of the tags was used in the amplicon pool, and iv) tag combinations where neither tag was used in the amplicon pool. For each amplicon library, proportions of total sequences within these categories were calculated. To compare tag-jump percentages between different treatments, unpaired t-tests were performed in R on percentage of sequences carrying new combinations of used tags within each amplicon pool (version 3.2.1, R Core Team, 2015).

## Results

### Effect of T4 DNA Polymerase blunt-ending and post-ligation PCR on tag-jump levels

To assess the amount of tag-jumps in the amplicon libraries created with the four library protocol treatments, within each library we calculated proportions of sequences carrying new combinations of used tags (tag-jumps). There was a significant effect of post-ligation PCR on tag-jump prevalence for both treatments with and without T4 DNA Polymerase blunt-ending. Libraries generated without T4 DNA Polymerase blunt-ending and including the post-ligation PCR step (−/+) had on average 22.7% of total sequences carrying new combinations of used tags (tag-jumps), which was significantly more than the Tagsteady protocol (−/−), which had on average 0.4% of total sequences carrying new combinations of used tags (unpaired t-test, p=0.03996) (Figure 3; Table S3). For the treatments with T4 DNA Polymerase blunt-ending, post-ligation PCR also resulted in a significantly higher amount of tag-jumps (+/+) (22.7%, unpaired t-test, P=0.0186) (Figure 3; Table S3).

**Figure 3.**
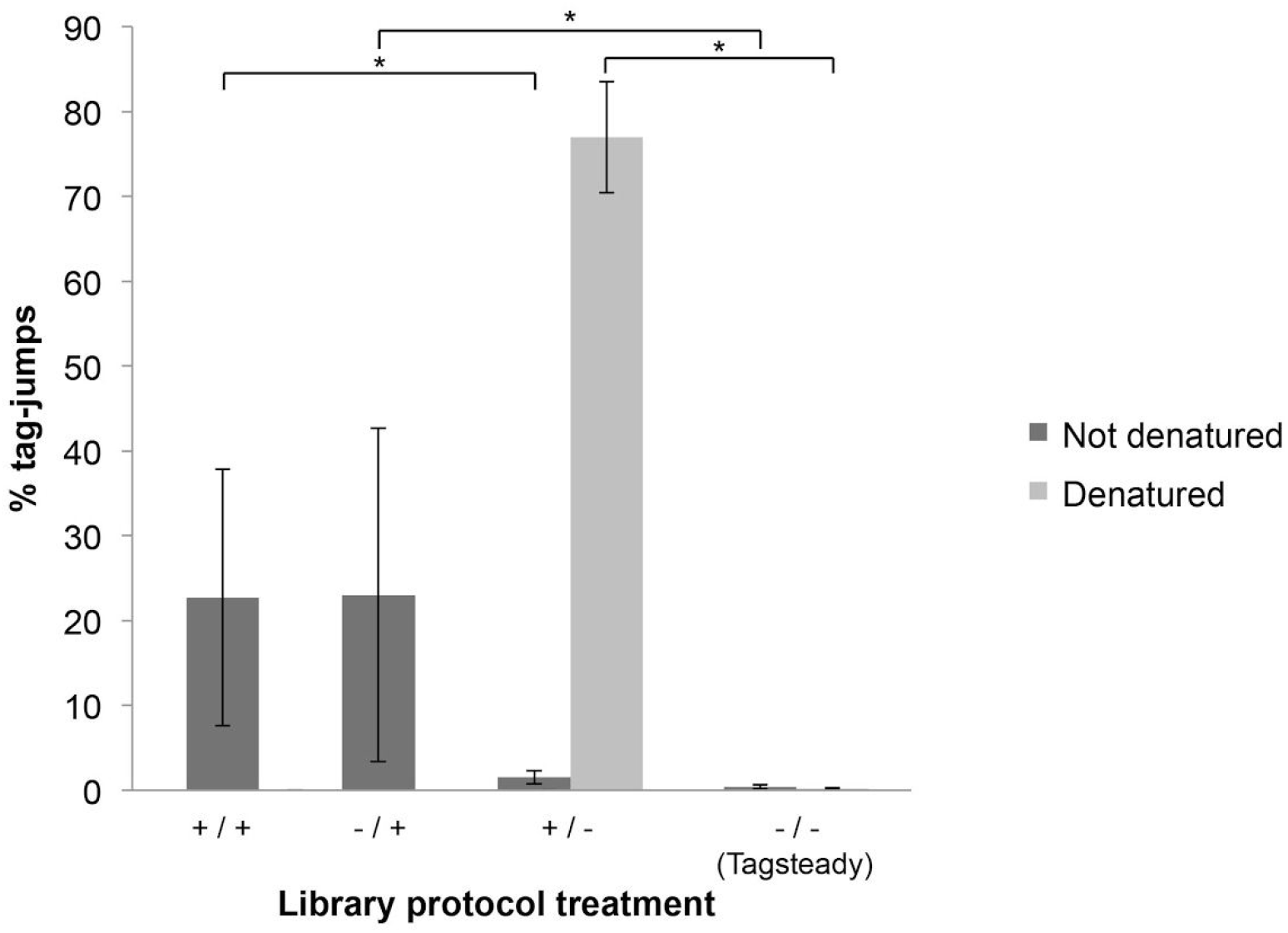
Average percentage of sequences carrying tag-jumps across amplicon pools built into Illumina libraries with four different library protocol treatments; +/+: T4 DNA polymerase blunt-ending and post-ligation PCR; −/+: no T4 DNA polymerase blunt-ending, with post-ligation PCR; +/−: T4 DNA polymerase blunt-ending and no post-ligation PCR; −/−: no T4 DNA polymerase blunt-ending and no post-ligation PCR (Tagsteady protocol) (n=6 for +/+, −/+, +/−, −/−). To mimic the effect of large amounts of single-stranded DNA generated in the metabarcoding PCR, aliquots of four of the amplicon pools were denatured and subsequently re-hybridized to form double-stranded DNA. These were then built into libraries with the −/− and +/− protocols (n=4 for d+/− and d−/−. Asterisks (*) denotes statistical significant difference between treatments (unpaired t-test, α=0.05).

Blunt-ending had little effect on the amount of tag-jumps (Figure 3). This led us to speculate that certain conditions in the metabarcoding PCR act as a prerequisite for tag-jumps. Specifically, if certain factors cause increased single-stranded DNA levels (e.g. difference in primer efficiency between forward and reverse primer or depletion of one primer during PCR amplification), this would cause more differently tagged single-stranded amplicons to form hybrids following pooling of amplicons. Consequently, this would result in more tag-jumps caused by the T4 DNA Polymerase.

### Effect of single-stranded amplicons on amount of tag-jumps

To test the hypothesis that T4 DNA Polymerase blunt-ending leads to increased tag-jump prevalence if it is carried out on amplicon pools with high amounts of hybrids of differently tagged single-stranded amplicons, we mimicked such a scenario by denaturing and re-hybridizing pools of tagged amplicons. Subsequently, we subjected them to the two library protocol treatments without post-ligation PCR: the library protocol treatment with T4 DNA Polymerase blunt-ending and without post-ligation PCR (+/−) and the Tagsteady library protocol (without T4 DNA Polymerase blunt-ending and without post-ligation PCR, −/−). Libraries built with T4 DNA Polymerase blunt-ending (+/−) had on average 76.99% (SD: 6.55) of total sequences carrying new combinations of used tags (tag-jumps) (Figure 3; Table S3). In contrast, libraries built with the Tagsteady protocol (−/−) had significantly lower amounts with an average of only 0.19% (SD: 0.13) of total sequences carrying new combinations of used tags (tag-jumps) (unpaired t-test, p-value = 0.0001692) (Figure 3; Table S3). This demonstrates that if a large amount of single-stranded DNA should be produced during the metabarcoding PCR, then tag-jump amounts can be high in the resulting data if end-repair is performed using blunt-ending. Moreover, it demonstrates that the Tagsteady protocol is able to cope with such a scenario.

### Validating the Tagsteady protocol

To validate the stability and robustness of the Tagsteady protocol, we used it to prepare libraries on 15 pools of twin-tagged amplicons. Across these amplicon libraries, the average number of sequences carrying new combinations of used tags (tag-jumps) was 0.25% (SD: 0.13) (Table 1). Overall, in this study, we carried out sequencing of 25 amplicon pools built into libraries using the Tagsteady protocol. The amplicon pools represented amplicons created with different metabarcoding primer sets and both denatured and undenatured amplicon pools. Across all of these amplicon libraries generated with the Tagsteady protocol, an average of 99.72% (SD: 0.17) of the sequences carried the expected tag combinations (range 99.22-99.96%) (Table S3).

**Table 1.** Validation of the Tagsteady library preparation protocol across 15 pools of twin-tagged amplicons. Range, median and average ±SD of percentage of total sequences assigned to four categories of tag combinations within each of the 15 amplicon pools. Only sequences with 100% match to primers and nucleotide tags are included.

| % total sequences within each of 15 pools of twin-tagged amplicons |  |  |  |
| --- | --- | --- | --- |
| Sequences carrying | Range (min - max) | Median | Average ± SD |
| Tag combinations where the tag pair was used in the amplicon pool | 99.418 - 99.882 | 99.769 | 99.746 ± 0.127 |
| New combination of tags used in the amplicon pool (**tag-jumps**) | 0.117 - 0.579 | 0.230 | 0.252 ± 0.127 |
| Tag combinations where only one of the tags was used in the amplicon pool | 0.0004 - 0.0030 | 0.001 | 0.0015 ± 0.0009 |
| Tag combinations where neither tag was used in the amplicon pool | 0.0000 - 0.0017 | 0.000 | 0.0002 ± 0.0004 |

The Tagsteady protocol routinely yielded a ligation efficiency high enough to provide ready-to-sequence libraries of 30 μl at 10-15 nM for amplicon pools in which the tagged primers did not have 5’ Ns added to increase complexity on the flow cell. For amplicon pools in which a number of random nucleotides (Ns) were added to the 5’ ends of primers to increase complexity on the flow cell, the ligation efficiency, and thereby the concentration of the final libraries, was lower (the lowest ca. 1 nM). The cause of the low ligation efficiency was most likely due to a high proportion of sequences with non-complementary ‘flapping’ ends that prohibited adapter ligation.

## Discussion

In this study, we presented and validated ‘Tagsteady’, a library preparation protocol for preparation of tagged amplicon pools for Illumina sequencing, and demonstrated that it can be used to practically eliminate tag-jumps. Further, we found that post-ligation PCR significantly increased tag-jump prevalence and that T4 DNA Polymerase blunt-ending caused high levels of tag-jumps if amplicon pools had high amounts of hybrids of differently tagged single-stranded amplicons (Fig. 3).

The use of the Tagsteady protocol yielded libraries with an average of 99.72 % (SD: 0.17) of sequences carrying the expected tag combinations (Table S3). All, or at least some, of these low levels of tag-jumps could be caused by sequencing errors in the tags (Coissac 2012) or tagged primer cross-contamination arising during synthesis and resuspension and during PCR reaction setup of tagged metabarcoding primers (see e.g. Kircher, Sawyer, and Meyer 2012). Consequently, we regard such low levels of background contamination to be unavoidable. Additionally, we note that it is possible to add internal controls to monitor the levels of such contamination through the inclusion of a few amplicons carrying unique forward and reverse tags, i.e. twin-tagged amplicons carrying tags not used on other amplicons in the amplicon pool.

When including post-ligation PCR, we found a high amount of sequences carrying tag-jumps (average of 22.7 and 23.0% for +/+ and −/+, respectively) compared to the findings of Schnell et al. 2015, who found an average of 3% of total sequences within amplicon pools to carry tag-jumps. This discrepancy between the two studies could be a result of different factors that can affect chimera levels in the post-ligation PCR, such as DNA template and primer concentration, PCR cycle number, and sequence similarity (e.g. Judo, Wedel, and Wilson 1998; Smyth et al. 2010). This range highlights the unreliability of a post-ligation PCR in metabarcoding studies and the importance of refining post-ligation PCR parameters to decrease the likelihood of chimera formation - or better yet, to completely omit the post-ligation PCR step.

Post-ligation PCR commonly serves two purposes, namely library enrichment and/or indexing of libraries (see e.g. Meyer and Kircher 2010). However, given that pools of tagged amplicons will contain high enough amounts of DNA to reach sufficient DNA concentration for sequencing and that indices can be added directly to the sequencing adapters, the post-ligation PCR step can be omitted when building libraries of pools of tagged amplicons, which we have demonstrated in the present protocol.

It should be noted that van Orsouw (2007) succeeded in using post-ligation in-solution PCR (required for the 454 platform) with no resulting tag-jumps on the 454 sequencing platform. However, this can be explained by their use of emulsion PCR, which prevents chimera formation, and therefore prevents tag-jumps. Although emulsion PCR could be used for post-ligation PCR in Illumina-based metabarcoding studies, it is not required given the aforementioned vast amounts of input material available for library preparation and the availability of PCR-free sequence adapters. Moreover, it adds to the cost and workload of library preparation.

As opposed to previous studies that demonstrated that T4 DNA Polymerase blunt-ending affects tag-jump prevalence (van Orsouw et al. 2007), we did not initially see an effect of blunt-ending (Figure 3). However, when subsequently building libraries on amplicon pools with high levels of hybrids of differently tagged single-stranded amplicons, we showed that the amount of single-stranded products in the metabarcoding PCR had been limiting our blunt-ending induced tag-jump amounts (Figure 3). This means that tag-jumps caused by blunt-ending can be reduced by refining PCR conditions to minimise the amount of single-stranded amplicons. For example, by keeping PCR cycle numbers at a minimum to avoid that it runs to completion. Other measures could be to ensure that forward and reverse primer concentrations are equimolar within each reaction and to ensure that their annealing temperatures are the same. All the amplicon pools used in this study were created using a relatively high concentration of metabarcoding primers and a limited number of cycles in the metabarcoding PCR to prevent the PCR reaction from plateauing (see Bohmann et al. 2018). This would have prevented depletion of available free primers, thereby minimizing the amount of single-stranded DNA and reducing blunt-ending induced tag-jump levels. Thus, it cannot be ruled out that with other metabarcoding PCR conditions we would have seen higher amounts of tag-jumps with the library protocol treatments that included T4 DNA Polymerase blunt-ending (+/− and+/+). However, here we show that the Tagsteady protocol enables practitioners to omit T4 DNA Polymerase blunt-ending and thereby circumvent the problem of tag-jumps caused by single-stranded amplicons.

Importantly, the Tagsteady protocol makes metabarcoding studies more cost-effective. First of all, a clear advantage of the virtually tag-jump free Tagsteady protocol, is that it allows practitioners to freely combine their tagged forward and reverse metabarcoding primers when carrying out metabarcoding PCRs. That is, where it was previously recommended to account for tag-jumps in the experimental setup through twin-tagging of amplicons, i.e. using of forward and reverse primers carrying matching tags, F1-R1, F2-R2, etc. (Schnell, Bohmann, and Gilbert 2015) a higher number of tag combinations can be made with the same number of tagged forward and reverse primers, which also means that fewer library preparations have to be made. Additionally, it will be less costly to buy primers as fewer tagged versions of each metabarcoding primer set have to be bought (Schnell, Bohmann, and Gilbert 2015). Further, as sequencing capacity will not be wasted on sequences carrying tag-jumps (here shown to be up to ca. 49 % of total sequences), the Tagsteady protocol makes sequencing more cost-effective. Lastly, to further reduce cost and workload, we designed the Tagsteady protocol as a single-tube library preparation protocol in which inter-reaction purification steps are omitted.

The Tagsteady protocol has a reasonably high ligation efficiency, routinely yielding a practical ligation efficiency of ~20-30% and ready-to-sequence amplicon libraries of 30 μl at 10-15 nM. It should, however, be noted that the ligation efficiency was greatly reduced when using the Tagsteady protocol to build libraries on amplicons to which random nucleotides (Ns) were added to the 5’ end of tags to increase complexity on the flow-cell. Therefore, when using the Tagsteady protocol, if nucleotides are added to the 5’ end of tags to increase complexity, then they should not be random (N’s). Rather, they should be ordered as specific nucleotides.

We applied the Tagsteady protocol to metabarcoding of eukaryotes in environmental samples. However, the protocol is applicable to amplicon library preparation of pools of other types of nucleotide tagged amplicons such as in 16S microbial studies (Caporaso et al. 2012), antibody sequencing (Menzel et al. 2014) and other gene targeted approaches where many sample DNA extracts are PCR amplified with 5’ tagged primers and pooled before Illumina library preparation. Further, while the Tagsteady protocol is designed for Illumina sequencing, it can be customised to other similar next-generation sequencing platforms, such as the Ion Torrent, by replacing Illumina adapters with appropriate sequencing platform specific adapters.

## Conclusion

The Tagsteady protocol is a fast, efficient and low-cost Illumina library preparation protocol for pools of 5’ nucleotide tagged amplicons that enables generation of metabarcoding data with correct assignments of sequences to samples, regardless of the extent of single-stranded DNA produced in the metabarcoding PCR.

We found that including T4 DNA Polymerase blunt-ending and/or post-ligation PCR in library preparation of pools of tagged amplicons can generate high amounts of false assignment of sequences to samples (tag-jumps), in this study up to ca. 49% of sequences within amplicon pools that had undergone post-ligation PCR carried tag-jumps. This highlights the fact that artefacts during library building for second generation sequencing can be detrimental to metabarcoding studies. We therefore encourage metabarcoding practitioners to avoid blunt-ending and post-ligation PCR during library preparation of amplicon pools consisting of 5’ tagged amplicons, e.g. through the use of the Tagsteady library preparation protocol. We advocate that commercial library kits designed for general shotgun sequencing libraries of genomic DNA are avoided for preparation of amplicon libraries as these will likely contain a T4 DNA Polymerase or a similar enzyme for end-repair, which could result in tag-jumps. If blunt-ending and post-ligation PCR cannot be avoided, for instance, if metabarcoding amplicons are sent for library preparation and sequencing at a commercial provider, we strongly advise to account for tag-jumps through the use of unique twin-tagged amplicons.

## Acknowledgements

We thank Lasse Vinner and Mette Juul Jacobsen at the Danish National High-Throughput DNA Sequencing Center for sequencing, discussions and invaluable guidance. We thank Tom Gilbert for support and discussions throughout the process.

## Funding

This work was supported by Independent Research Fund Denmark, DFF grant 5051-00140 (KB), Innovation fund Denmark ‘Foodtranscriptomics’ (50557100-1118631003) and Carlsberg Foundation Semper Ardens ‘Archives’ (CF18-1110) (CC).

## Author contributions

CC and KB conceived the ideas, performed experiments, analyzed the data and wrote the paper.

## Competing interests

The authors declare no competing financial interests.

## Supplementary information

Additional Information can be found in the online version of this article:

Appendix S1: Adapter design, synthesis and preparation

Appendix S2: Table S2. Overview of experimental setup

Appendix S3: Supplementary results

Table S3a. Overview of proportion of expected and unexpected categories of tag combinations following different library protocol treatments.

## Data accessibility

Sequence data and primer and tag information will be uploaded to the Dryad Digital Repository, DOI:xxxxxxx upon acceptance of the manuscript.

## References

Alberdi, Antton, Ostaizka Aizpurua, Kristine Bohmann, Shyam Gopalakrishnan, Christina Lynggaard, Martin Nielsen, and Marcus Thomas Pius Gilbert. 2018. “Promises and Pitfalls of Using High-Throughput Sequencing for Diet Analysis.” Molecular Ecology Resources, October. https://doi.org/10.1111/1755-0998.12960.

Apothéloz-Perret-Gentil, Laure, Arielle Cordonier, François Straub, Jennifer Iseli, Philippe Esling, and Jan Pawlowski. 2017. “Taxonomy-Free Molecular Diatom Index for High-Throughput eDNA Biomonitoring.” Molecular Ecology Resources 17 (6): 1231–42.

Bentley, David R., Shankar Balasubramanian, Harold P. Swerdlow, Geoffrey P. Smith, John Milton, Clive G. Brown, Kevin P. Hall, et al. 2008. “Accurate Whole Human Genome Sequencing Using Reversible Terminator Chemistry.” Nature 456 (7218): 53–59.

Binladen, Jonas, M. Thomas P. Gilbert, Jonathan P. Bollback, Frank Panitz, Christian Bendixen, Rasmus Nielsen, and Eske Willerslev. 2007. “The Use of Coded PCR Primers Enables High-Throughput Sequencing of Multiple Homolog Amplification Products by 454 Parallel Sequencing.” PloS One 2 (2): e197.

Blaalid, Rakel, Surendra Kumar, R. Henrik Nilsson, Kessy Abarenkov, P. M. Kirk, and H. Kauserud. 2013. “ITS 1 versus ITS 2 as DNA Metabarcodes for Fungi.” Molecular Ecology Resources 13 (2): 218–24.

Blaalid, R., M. L. Davey, H. Kauserud, and T. Carlsen. 2014. “Arctic Root-associated Fungal Community Composition Reflects Environmental Filtering.” Molecular. https://onlinelibrary.wiley.com/doi/abs/10.1111/mec.12622.

Bohmann, Kristine, Alice Evans, M. Thomas P. Gilbert, Gary R. Carvalho, Simon Creer, Michael Knapp, Douglas W. Yu, and Mark de Bruyn. 2014. “Environmental DNA for Wildlife Biology and Biodiversity Monitoring.” Trends in Ecology & Evolution 29 (6): 358–67.

Bohmann, Kristine, Shyam Gopalakrishnan, Martin Nielsen, Luisa Dos Santos Bay Nielsen, Gareth Jones, Daniel G. Streicker, and M. Thomas P. Gilbert. 2018. “Using DNA Metabarcoding for Simultaneous Inference of Common Vampire Bat Diet and Population Structure.” Molecular Ecology Resources, April. https://doi.org/10.1111/1755-0998.12891.

Bohmann, Kristine, Ara Monadjem, Christina Lehmkuhl Noer, Morten Rasmussen, Matt R. K. Zeale, Elizabeth Clare, Gareth Jones, Eske Willerslev, and M. Thomas P. Gilbert. 2011. “Molecular Diet Analysis of Two African Free-Tailed Bats (molossidae) Using High Throughput Sequencing.” PloS One 6 (6): e21441.

Botnen, Synnøve, Unni Vik, Tor Carlsen, Pernille B. Eidesen, Marie L. Davey, and Håvard Kauserud. 2014. “Low Host Specificity of Root-Associated Fungi at an Arctic Site.” Molecular Ecology 23 (4): 975–85.

Caporaso, J. Gregory, Christian L. Lauber, William A. Walters, Donna Berg-Lyons, James Huntley, Noah Fierer, Sarah M. Owens, et al. 2012. “Ultra-High-Throughput Microbial Community Analysis on the Illumina HiSeq and MiSeq Platforms.” The ISME Journal 6 (8): 1621–24.

Carew, Melissa E., Vincent J. Pettigrove, Leon Metzeling, and Ary A. Hoffmann. 2013. “Environmental Monitoring Using next Generation Sequencing: Rapid Identification of Macroinvertebrate Bioindicator Species.” Frontiers in Zoology 10 (1): 45.

Carøe, Christian, Shyam Gopalakrishnan, Lasse Vinner, Sarah S. T. Mak, Mikkel Holger S. Sinding, José A. Samaniego, Nathan Wales, Thomas Sicheritz-Pontén, and M. Thomas P. Gilbert. 2018. “Single-Tube Library Preparation for Degraded DNA.” Methods in Ecology and Evolution / British Ecological Society 9 (2): 410–19.

Clarke, Laurence J., Julien Soubrier, Laura S. Weyrich, and Alan Cooper. 2014. “Environmental Metabarcodes for Insects: In Silico PCR Reveals Potential for Taxonomic Bias.” Molecular Ecology Resources 14 (6): 1160–70.

Coissac, Eric. 2012. “OligoTag: A Program for Designing Sets of Tags for next-Generation Sequencing of Multiplexed Samples.” Methods in Molecular Biology 888: 13–31.

Davey, Marie L., Rune Heimdal, Mikael Ohlson, and Håvard Kauserud. 2013. “Host- and Tissue-Specificity of Moss-Associated Galerina and Mycena Determined from Amplicon Pyrosequencing Data.” Fungal Ecology 6 (3): 179–86.

Davey, Marie L., Håvard Kauserud, and Mikael Ohlson. 2014. “Forestry Impacts on the Hidden Fungal Biodiversity Associated with Bryophytes.” FEMS Microbiology Ecology 90 (1): 313–25.

De Barba, M., C. Miquel, F. Boyer, C. Mercier, D. Rioux, E. Coissac, and P. Taberlet. 2014. “DNA Metabarcoding Multiplexing and Validation of Data Accuracy for Diet Assessment: Application to Omnivorous Diet.” Molecular Ecology Resources 14 (2): 306–23.

Elbrecht, Vasco, Pierre Taberlet, Tony Dejean, Alice Valentini, Philippe Usseglio-Polatera, Jean-Nicolas Beisel, Eric Coissac, Frederic Boyer, and Florian Leese. 2016. “Testing the Potential of a Ribosomal 16S Marker for DNA Metabarcoding of Insects.” PeerJ 4 (April): e1966.

Elbrecht, Vasco, Ecaterina Edith Vamos, Kristian Meissner, Jukka Aroviita, and Florian Leese. 2017. “Assessing Strengths and Weaknesses of DNA Metabarcoding-Based Macroinvertebrate Identification for Routine Stream Monitoring.” Methods in Ecology and Evolution / British Ecological Society 8 (10): 1265–75.

Esling, Philippe, Franck Lejzerowicz, and Jan Pawlowski. 2015. “Accurate Multiplexing and Filtering for High-Throughput Amplicon-Sequencing.” Nucleic Acids Research 43 (5): 2513–24.

Faircloth, B. C., and T. C. Glenn. 2014. “Protocol: Preparation of an AMPure XP Substitute (AKA Serapure). Doi: 10.6079.” J9MW2F26.

Geller, John, C. Meyer, M. Parker, and H. Hawk. 2013. “Redesign of PCR Primers for Mitochondrial Cytochrome c Oxidase Subunit I for Marine Invertebrates and Application in All-Taxa Biotic Surveys.” Molecular Ecology Resources 13 (5): 851–61.

Hope, Paul R., Kristine Bohmann, M. Thomas P. Gilbert, Marie Lisandra Zepeda-Mendoza, Orly Razgour, and Gareth Jones. 2014. “Second Generation Sequencing and Morphological Faecal Analysis Reveal Unexpected Foraging Behaviour by Myotis Nattereri (Chiroptera, Vespertilionidae) in Winter.” Frontiers in Zoology 11 (1): 39.

Judo, M. S., A. B. Wedel, and C. Wilson. 1998. “Stimulation and Suppression of PCR-Mediated Recombination.” Nucleic Acids Research 26 (7): 1819–25.

Kircher, Martin, Susanna Sawyer, and Matthias Meyer. 2012. “Double Indexing Overcomes Inaccuracies in Multiplex Sequencing on the Illumina Platform.” Nucleic Acids Research 40 (1): e3.

Kozarewa, Iwanka, Zemin Ning, Michael A. Quail, Mandy J. Sanders, Matthew Berriman, and Daniel J. Turner. 2009. “Amplification-Free Illumina Sequencing-Library Preparation Facilitates Improved Mapping and Assembly of (G+C)-Biased Genomes.” Nature Methods 6 (4): 291–95.

Leray, Matthieu, Joy Y. Yang, Christopher P. Meyer, Suzanne C. Mills, Natalia Agudelo, Vincent Ranwez, Joel T. Boehm, and Ryuji J. Machida. 2013. “A New Versatile Primer Set Targeting a Short Fragment of the Mitochondrial COI Region for Metabarcoding Metazoan Diversity: Application for Characterizing Coral Reef Fish Gut Contents.” Frontiers in Zoology 10 (1): 34.

Lindner, Daniel L., Tor Carlsen, R. Henrik Nilsson, Marie Davey, Trond Schumacher, and Håvard Kauserud. 2013. “Employing 454 Amplicon Pyrosequencing to Reveal Intragenomic Divergence in the Internal Transcribed Spacer rDNA Region in Fungi.” Ecology and Evolution 3 (6): 1751–64.

Mak, S. S. T., S. Gopalakrishnan, C. Carøe, and C. Geng. 2017. “Comparative Performance of the BGISEQ-500 vs Illumina HiSeq2500 Sequencing Platforms for Palaeogenomic Sequencing.” https://academic.oup.com/gigascience/article-abstract/6/8/1/3958782.

Margulies, Marcel, Michael Egholm, William E. Altman, Said Attiya, Joel S. Bader, Lisa A. Bemben, Jan Berka, et al. 2005. “Genome Sequencing in Microfabricated High-Density Picolitre Reactors.” Nature 437 (7057): 376–80.

Menzel, Ulrike, Victor Greiff, Tarik A. Khan, Ulrike Haessler, Ina Hellmann, Simon Friedensohn, Skylar C. Cook, Mark Pogson, and Sai T. Reddy. 2014. “Comprehensive Evaluation and Optimization of Amplicon Library Preparation Methods for High-Throughput Antibody Sequencing.” PloS One 9 (5): e96727.

Meyer, Matthias, and Martin Kircher. 2010. “Illumina Sequencing Library Preparation for Highly Multiplexed Target Capture and Sequencing.” Cold Spring Harbor Protocols 2010 (6): db.prot5448.

Neiman, Mårten, Simon Sundling, Henrik Grönberg, Per Hall, Kamila Czene, Johan Lindberg, and Daniel Klevebring. 2012. “Library Preparation and Multiplex Capture for Massive Parallel Sequencing Applications Made Efficient and Easy.” PloS One 7 (11): e48616.

Orsouw, Nathalie J. van, René C. J. Hogers, Antoine Janssen, Feyruz Yalcin, Sandor Snoeijers, Esther Verstege, Harrie Schneiders, et al. 2007. “Complexity Reduction of Polymorphic Sequences (CRoPS^TM^): A Novel Approach for Large-Scale Polymorphism Discovery in Complex Genomes.” PloS One 2 (11): e1172.

Palkopoulou, Eleftheria, Mateusz Baca, Natalia I. Abramson, Mikhail Sablin, Paweł Socha, Adam Nadachowski, Stefan Prost, et al. 2016. “Synchronous Genetic Turnovers across Western Eurasia in Late Pleistocene Collared Lemmings.” Global Change Biology 22 (5): 1710–21.

Quéméré, Erwan, Fabrice Hibert, Christian Miquel, Emeline Lhuillier, Emmanuel Rasolondraibe, Julie Champeau, Clément Rabarivola, et al. 2013. “A DNA Metabarcoding Study of a Primate Dietary Diversity and Plasticity across Its Entire Fragmented Range.” PloS One 8 (3): e58971.

Rittié, Laure, and Bernard Perbal. 2008. “Enzymes Used in Molecular Biology: A Useful Guide.” Journal of Cell Communication and Signaling 2 (1-2): 25–45.

Rohland, Nadin, and David Reich. 2012. “Cost-Effective, High-Throughput DNA Sequencing Libraries for Multiplexed Target Capture.” Genome Research 22 (5): 939–46.

Schnell, Ida Bærholm, Kristine Bohmann, and M. Thomas P. Gilbert. 2015. “Tag Jumps Illuminated--Reducing Sequence-to-Sample Misidentifications in Metabarcoding Studies.” Molecular Ecology Resources 15 (6): 1289–1303.

Schubert, Mikkel, Stinus Lindgreen, and Ludovic Orlando. 2016. “AdapterRemoval v2: Rapid Adapter Trimming, Identification, and Read Merging.” BMC Research Notes 9 (February): 88.

Shehzad, Wasim, Tiayyba Riaz, Muhammad A. Nawaz, Christian Miquel, Carole Poillot, Safdar A. Shah, Francois Pompanon, Eric Coissac, and Pierre Taberlet. 2012. “Carnivore Diet Analysis Based on next-Generation Sequencing: Application to the Leopard Cat (Prionailurus Bengalensis) in Pakistan.” Molecular Ecology 21 (8): 1951–65.

Singer, Greg, Nicole A. Fahner, Joshua Barnes, Avery McCarthy, and Mehrdad Hajibabaei. 2019. “Comprehensive Biodiversity Analysis via Ultra-Deep Patterned Flow Cell Technology: A Case Study of eDNA Metabarcoding Seawater.” bioRxiv. https://doi.org/10.1101/515890.

Sinha, Rahul, Geoff Stanley, Gunsagar Singh Gulati, Camille Ezran, Kyle Joseph Travaglini, Eric Wei, Charles Kwok Fai Chan, et al. 2017. “Index Switching Causes ‘Spreading-Of-Signal’ Among Multiplexed Samples In Illumina HiSeq 4000 DNA Sequencing.” bioRxiv. https://doi.org/10.1101/125724.

Smyth, R. P., T. E. Schlub, A. Grimm, V. Venturi, A. Chopra, S. Mallal, M. P. Davenport, and J. Mak. 2010. “Reducing Chimera Formation during PCR Amplification to Ensure Accurate Genotyping.” Gene 469 (1-2): 45–51.

Stoeck, Thorsten, Hongbo Pan, Verena Dully, Dominik Forster, and Thorsten Jung. 2018. “Towards an eDNA Metabarcode-Based Performance Indicator for Full-Scale Municipal Wastewater Treatment Plants.” Water Research 144 (November): 322–31.

Taberlet, Pierre, Eric Coissac, François Pompanon, Christian Brochmann, and Eske Willerslev. 2012. “Towards next-Generation Biodiversity Assessment Using DNA Metabarcoding.” Molecular Ecology 21 (8): 2045–50.

Taylor, P. G. 1996. “Reproducibility of Ancient DNA Sequences from Extinct Pleistocene Fauna.” Molecular Biology and Evolution 13 (1): 283–85.

Valentini, Alice, Christian Miquel, Muhammad Ali Nawaz, Eva Bellemain, Eric Coissac, François Pompanon, Ludovic Gielly, et al. 2009. “New Perspectives in Diet Analysis Based on DNA Barcoding and Parallel Pyrosequencing: The trnL Approach.” Molecular Ecology Resources 9 (1): 51–60.

Zepeda-Mendoza, Marie Lisandra, Kristine Bohmann, Aldo Carmona Baez, and M. Thomas P. Gilbert. 2016. “DAMe: A Toolkit for the Initial Processing of Datasets with PCR Replicates of Double-Tagged Amplicons for DNA Metabarcoding Analyses.” BMC Research Notes 9 (May): 255.

Zheng, Zongli, Abdolreza Advani, Ojar Melefors, Steve Glavas, Henrik Nordström, Weimin Ye, Lars Engstrand, and Anders F. Andersson. 2010. “Titration-Free Massively Parallel Pyrosequencing Using Trace Amounts of Starting Material.” Nucleic Acids Research 38 (13): e137.

