## Supplemental information for "Tagsteady: a metabarcoding library preparation protocol to avoid false assignment of sequences to samples"

C. Carøe and K. Bohmann

Appendix S1: Adapter design, synthesis and preparation
Appendix S2: Table S2 Overview of experimental setup
Appendix S3: Table S3 Supplementary results

**Appendix S1: Adapter design, synthesis and preparation**

Adapter oligos were similar to Illuminas full-length PCR-free Truseq adapters (support.illumina.com) albeit with indices of seven nucleotides in length and with a minimum of three differences between any two indices (Table S1). Adapter oligos were ordered from Integrated DNA Technologies (IDT inc. Iowa, US), synthesized at 1 µM scale, purified with PAGE purification and shipped dry. Oligos were dissolved to 500 µM in TET buffer (1 mM EDTA, 10 mM tris-HCl and 0.05% Tween-20, Sigma-Aldrich, P9416-50ML), incubated at room temperature for 10 minutes and mixed by mild vortexing. 20 µL (500 µM) P5 was mixed with 20 µL (500 µM) P7 and 5 µl water and 5 µl oligo hybridization buffer (500 mM NaCl, 10 mM Tris-HCl, 1 mM EDTA) were added. This was mixed well and incubated in a heat block for 10 sec at 95°C before ramping down to 12°C with a rate of 0.1°C per second. Finally, adapters were diluted with EBT (10 mM Tris-HCl, 0.05% Tween-20)to a working solution of 20 µM. Adapters were aliquoted into 20 µl aliquots to prevent excessive freeze-thaw cycles and stored at -20°C. A total of 8 adapter mixes were prepared with unique matching index combination, (i.e. 1-1, 2-2, etc.).

**References**

Illumina Adapter Sequences (1000000002694 v08) Posted 10/02/2018: https://support.illumina.com/downloads/illumina-adapter-sequences-document-1000000002694.html

### Table S1.

* indicates Phosphothioate bond, lowercase indicates index, [Phosphate] indicates 5’ phosphate. 501-508 are P5 oligos, while 701-708 are P7 oligos.

| Name | Oligo sequence (5’-3’) |
|---|---|
| 501 | AATGATACGGCGACCACCGAGATCTACACcctgcgaACACTCTTTCCCTACACGACGCTCTTCCGATC*T |
| 502 | AATGATACGGCGACCACCGAGATCTACACtgcagagACACTCTTTCCCTACACGACGCTCTTCCGATC*T |
| 503 | AATGATACGGCGACCACCGAGATCTACACacctaggACACTCTTTCCCTACACGACGCTCTTCCGATC*T |
| 504 | AATGATACGGCGACCACCGAGATCTACACttgatccACACTCTTTCCCTACACGACGCTCTTCCGATC*T |
| 505 | AATGATACGGCGACCACCGAGATCTACACatcttgcACACTCTTTCCCTACACGACGCTCTTCCGATC*T |
| 506 | AATGATACGGCGACCACCGAGATCTACACtctccatACACTCTTTCCCTACACGACGCTCTTCCGATC*T |
| 507 | AATGATACGGCGACCACCGAGATCTACACcatcgagACACTCTTTCCCTACACGACGCTCTTCCGATC*T |
| 508 | AATGATACGGCGACCACCGAGATCTACACttcgagcACACTCTTTCCCTACACGACGCTCTTCCGATC*T |
| 701 | [Phosphate]GATCGGAAGAGCACACGTCTGAACTCCAGTCACcctgcgaATCTCGTATGCCGTCTTCTGCTTG |
| 702 | [Phosphate]GATCGGAAGAGCACACGTCTGAACTCCAGTCACtgcagagATCTCGTATGCCGTCTTCTGCTTG |
| 703 | [Phosphate]GATCGGAAGAGCACACGTCTGAACTCCAGTCACacctaggATCTCGTATGCCGTCTTCTGCTTG |
| 704 | [Phosphate]GATCGGAAGAGCACACGTCTGAACTCCAGTCACttgatccATCTCGTATGCCGTCTTCTGCTTG |
| 705 | [Phosphate]GATCGGAAGAGCACACGTCTGAACTCCAGTCACatcttgcATCTCGTATGCCGTCTTCTGCTTG |
| 706 | [Phosphate]GATCGGAAGAGCACACGTCTGAACTCCAGTCACtctccatATCTCGTATGCCGTCTTCTGCTTG |
| 707 | [Phosphate]GATCGGAAGAGCACACGTCTGAACTCCAGTCACcatcgagATCTCGTATGCCGTCTTCTGCTTG |

| | |
|---|---|
| 708 | [Phosphate]GATCGGAAGAGCACACGTCTGAACTCCAGTCACttcgagcATCTCGTATGCCGTCTTCTGCTTG |

### Appendix S2. Overview of experimental set-up

**Table S2.** Overview of experimental set-up. Four library preparation protocol treatments were carried out on six pools of tagged amplicons. The four library protocols represent combinations with and without T4 DNA Polymerase blunt-ending and with and without post-ligation PCR. Further, two of the library protocol treatments were carried out on denatured and re-hybridized aliquots of four of the amplicon pools to assess whether the Tagsteady protocol can withstand high proportions of single-stranded amplicons.

| **Metabarcoding marker** | **Amplicon pool** | **Blunt-ending using T4 DNA polymerase in the end-repair step** | **12 cycles of post-library amplification** | **Library protocol treatment** |
|---|---|---|---|---|
| Metazoan COI | COI_1 | + | + | +/+ |
| | COI_1 | + | - | +/- |
| | COI_1 | - | - | -/- |
| | COI_1 | - | + | -/+ |
| | COI_2 | + | + | +/+ |
| | COI_2 | + | - | +/- |
| | COI_2 | - | - | -/- |
| | COI_2 | - | + | -/+ |
| Insect 16sRNA | 16sIns_1 | + | + | +/+ |
| | 16sIns_1 | + | - | +/- |
| | 16sIns_1 | - | - | -/- |
| | 16sIns_1 | - | + | -/+ |
| | 16sIns_2 | + | + | +/+ |
| | 16sIns_2 | + | - | +/- |
| | 16sIns_2 | - | - | -/- |
| | 16sIns_2 | - | + | -/+ |
| Mammal 16sRNA | 16sMam_1 | + | + | +/+ |
| | 16sMam_1 | + | - | +/- |
| | 16sMam_1 | - | - | -/- |
| | 16sMam_1 | - | + | -/+ |
| | 16sMam_2 | + | + | +/+ |
| | 16sMam_2 | + | - | +/- |
| | 16sMam_2 | - | - | -/- |
| | 16sMam_2 | - | + | -/+ |
| Negative controls | Neg_1 & _2 | + | - | +/- |
| | Neg__3 & _4 | - | - | -/- |
| | Neg_5 & _6 | + | + | +/+ |
| | Neg_7 & _8 | - | + | -/+ |
| Insect 16sRNA | 16sIns_1 | + | - | D+/- |
| | 16sIns_1 | - | - | D-/- |
| | 16sIns_2 | + | - | D+/- |
| | 16sIns_2 | - | - | D-/- |
| Mammal 16sRNA | 16sMam_1 | + | - | D+/- |
| | 16sMam_1 | - | - | D-/- |
| | 16sMam_2 | + | - | D+/- |
| | 16sMam_2 | - | - | D-/- |

**Appendix S3: Supplementary results**

**Table S3.** Overview of proportion of expected and unexpected categories of tag combinations following different library protocol treatments. Four library preparation protocols were carried out on six pools of tagged amplicons representing combinations with and without T4 DNA Polymerase blunt-ending and with and without post-ligation PCR. Further, two of the library protocol treatments were carried out on denatured and re-hybridized aliquots of four of the amplicon pools to assess whether the Tagsteady protocol can withstand high proportions of single-stranded amplicons.

| Library protocol treatment | Amplicon pool | Tag combinations where the tag pair was used | Tag combinations where both tags were used but not in this combination (tag jumps) | Tag combinations where only one of the tags was used | Tag combinations where neither tag was used |
|---|---|---|---|---|---|
| | | Number sequences (%) | | | |
| +T4 DNA polymerase / + library PCR | 16sMam_1_+/+ | 506,762 (58.32 %) | 361,886 (41.65 %) | 285 (0.03 %) | 0 (0 %) |
| | 16sMam_2_+/+ | 511,563 (59.75 %) | 344,407 (40.23 %) | 141 (0.02 %) | 0 (0 %) |
| | 16sIns_1_+/+ | 1052418 (91.69 %) | 95,328 (8.31 %) | 11 (0.00 %) | 0 (0.00 %) |
| | 16sIns_2_+/+ | 1,017,194 (92.98 %) | 76,800 (7.02 %) | 5 (0.00 %) | 0 (0.00%) |
| | COI_metaz_1_+/+ | 562,258 (78.07 %) | 157,936 (21.93 %) | 35 (0.00 %) | 0 (0.00 %) |
| | COI_metaz_2_+/+ | 584,911 (82.76 %) | 121,775 (17.23 %) | 98 (0.01 %) | 1 (0.00 %) |
| - T4 DNA polymerase / + library PCR | 16sMam_1_-/+ | 508,353 (50.74 %) | 493,289 (49.23 %) | 291 (0.03 %) | 13 (0.00 %) |
| | 16sMam_2_-/+ | 396,246 (52.78 %) | 354,427 (47.21 %) | 98 (0.01 %) | 1 (0.00 %) |
| | 16sIns_1_-/+ | 705, 427 (91.76 %) | 63,352 (8.24 %) | 11 (0.00 %) | 0 (0.00 %) |
| | 16sIns_2_-/+ | 948,458 (90.63 %) | 98,018 (9.37 %) | 5 (0.00 %) | 0 (0.00 %) |
| | COI_metaz_1_-/+ | 789,291 (86.22 %) | 126,140 (13.78 %) | 37 (0.00 %) | 0 (0.00 %) |
| | COI_metaz_2_-/+ | 989,957 (89.78 %) | 112,547 (10.21 %) | 115 (0.01 %) | 1 (0.00 %) |
| + T4 DNA polymerase / - library PCR | 16sMam_1_+/- | 1,658,806 (99.01 %) | 16,089 (0.96 %) | 423 (0.03 %) | 7 (0.00 %) |
| | 16sMam_2_+/- | 1,746,063 (99.6 %) | 6,639 (0.38 %) | 365 (0.02 %) | 7 (0.00 %) |
| | 16sIns_1_+/- | 1,025,368 (97.36 %) | 27,779 (2.64 %) | 9 (0.00 %) | 0 (0.00 %) |
| | 16sIns_2_+/- | 881,087 (97.96 %) | 18,293 (2.03 %) | 10 (0.00 %) | 0 (0.00 %) |
| | COI_metaz_1_+/- | 730,822 (98.44 %) | 11,583 (1.56 %) | 27 (0.00 %) | 0 (0.00 %) |
| | COI_metaz_2_+/- | 738,838 (98.47 %) | 11,366 (1.51 %) | 121 (0.02 %) | 0 (0.00 %) |
| - T4 DNA polymerase / - library PCR | 16sMam_1_-/- | 1,940,874 (99.49 %) | 9,748 (0.5 %) | 285 (0.01 %) | 6 (0.00 %) |
| | 16sMam_2_-/- | 1,577,004 (99.89 %) | 1,523 (0.1 %) | 213 (0.01 %) | 2 (0.00%) |
| | 16sIns_1_-/- | 1,030,345 (99.76 %) | 2,486 (0.24 %) | 8 (0.00 %) | 0 (0.00 %) |
| | 16sIns_2_-/- | 815,554 (99.76 %) | 1,948 (0.24 %) | 6 (0.00 %) | 0 (0.00 %) |

| | | | | | |
|---|---|---|---|---|---|
| | COI_metaz_1_-/- | 636,263 (99.22 %) | 4,978 (0.78 %) | 25 (0.00 %) | 0 (0.00 %) |
| | COI_metaz_2_-/- | 710,998 (99.47 %) | 3,737 (0.52 %) | 77 (0.01 %) | 0 (0.00 %) |
| + T4 DNA polymerase / - library PCR on denatured amplicon pool | 16sMam_1_D+/- | 247,663 (30.76 %) | 556,820 (69.17 %) | 191 (0.02 %) | 350 (0.04 %) |
| | 16sMam_2_D+/- | 112,260 (25.86 %) | 321,752 (74.13 %) | 47 (0.01 %) | 0 (0.00 %) |
| | 16sIns_1_D+/- | 49,903 (18.72 %) | 216,581 (81.26 %) | 23 (0.01 %) | 27 (0.01 %) |
| | 16sIns_2_D+/- | 217,188 (16.58 %) | 1,092,409 (83.41 %) | 38 (0 %) | 0 (0.00%) |
| - T4 DNA polymerase / - library PCR on denatured amplicon pool | 16sMam_1_D-/- | 1,774,615 (99.71 %) | 5,169 (0.29 %) | 74 (0.00 %) | 5 (0.00 %) |
| | 16sMam_2_D-/- | 2,176,298 (99.70 %) | 6,482 (0.30 %) | 85 (0.00 %) | 2 (0.00 %) |
| | 16sIns1_D-/- | 2,544,537 (99.96 %) | 998 (0.04 %) | 6 (0.00 %) | 0 (0.00 %) |
| | 16sIns2_D-/- | 2,243,003 (99.88 %) | 2,656 (0.12 %) | 5 (0.00 %) | 0 (0.00 %) |
